# Coarse-grained molecular simulations of the binding of the SARS-CoV-2 spike protein RBD to the ACE2 receptor

**DOI:** 10.1101/2020.05.07.083212

**Authors:** David De Sancho, José A. Gavira, Raul Pérez-Jiménez

**Author notes:** Corresponding author Email address (David De Sancho).

## Abstract

Since it was first observed, the COVID-19 pandemic has created a global emergency for national health systems due to millions of confirmed cases and hundreds of thousands of deaths. At a molecular level, the bottleneck for the infection is the binding of the receptor binding domain (RBD) of the viral spike protein to ACE2, an enzyme exposed on human cell membranes. Several experimental structures of the ACE2:RBD complex have been made available, however they offer only a static description of the arrangements of the molecules in either the free or bound states. In order to gain a dynamic description of the binding process that is key to infection, we use molecular simulations with a coarse grained model of the RBD and ACE2. We find that binding occurs in an all-or-none way, without intermediates, and that even in the bound state, the RBD exhibits a considerably dynamic behaviour. From short equilibrium simulations started in the unbound state we provide snapshots that result in a tentative mechanism of binding. Our findings may be important for the development of drug discovery strategies that target the RBD.

## 1. Introduction

The spillover of the severe acute respiratory syndrome coronavirus 2 (SARS-CoV-2) to humans has resulted in the coronavirus disease 2019 (or COVID-19) pandemic [1]. The number of cases around the planet has skyrocketed in many countries, causing major disruptions and hundreds of thousands of deaths around the world. This crisis has spurred titanic efforts from the scientific community in order to develop therapies to combat the disease and prevent future outbreaks. In particular, structural biologists have been quick to resolve the 3D coordinates of a few of the viral proteins, including among others its main protease [2], the Nsp15 endoribonuclease [3], and the spike protein [4, 5, 6, 7, 8]. These efforts provide invaluable detail of the molecular machinery involved in the viral infection cycle, which is essential for the development of effective drugs.

Among the therapeutic strategies aiming to stop the pandemic, much work is being devoted to target the interaction between the trimeric spike protein (or protein S) to the host receptor, the angiotensin-converting enzyme 2 (ACE2) [9, 10]. The binding between both proteins precedes membrane fusion and may be the Achilles heel of the virus. The spike first engages with the host cell when it interacts with ACE2 via the very mobile receptor binding domain (RBD) of one of its monomers in its “up” state [4]. The interaction between the RBD and ACE2 buries an extended surface of ∼1700 Å ^2^ involving a coiled coil formed by two long helices of ACE2 and a loosely packed binding motif in the RBD (see Figure 1 A). The structure for the SARS-CoV-2 RBD bound to ACE2 bears close resemblance to that of the same complex for the previous SARS-CoV virus [11]. However, there is evidence that the greater infectivity of the new virus may be determined at least in part by the ∼10-20-fold higher binding affinity for ACE2 [4] (although estimates differ among groups, see Supplementary Information Fig. S1).

**Figure 1:**
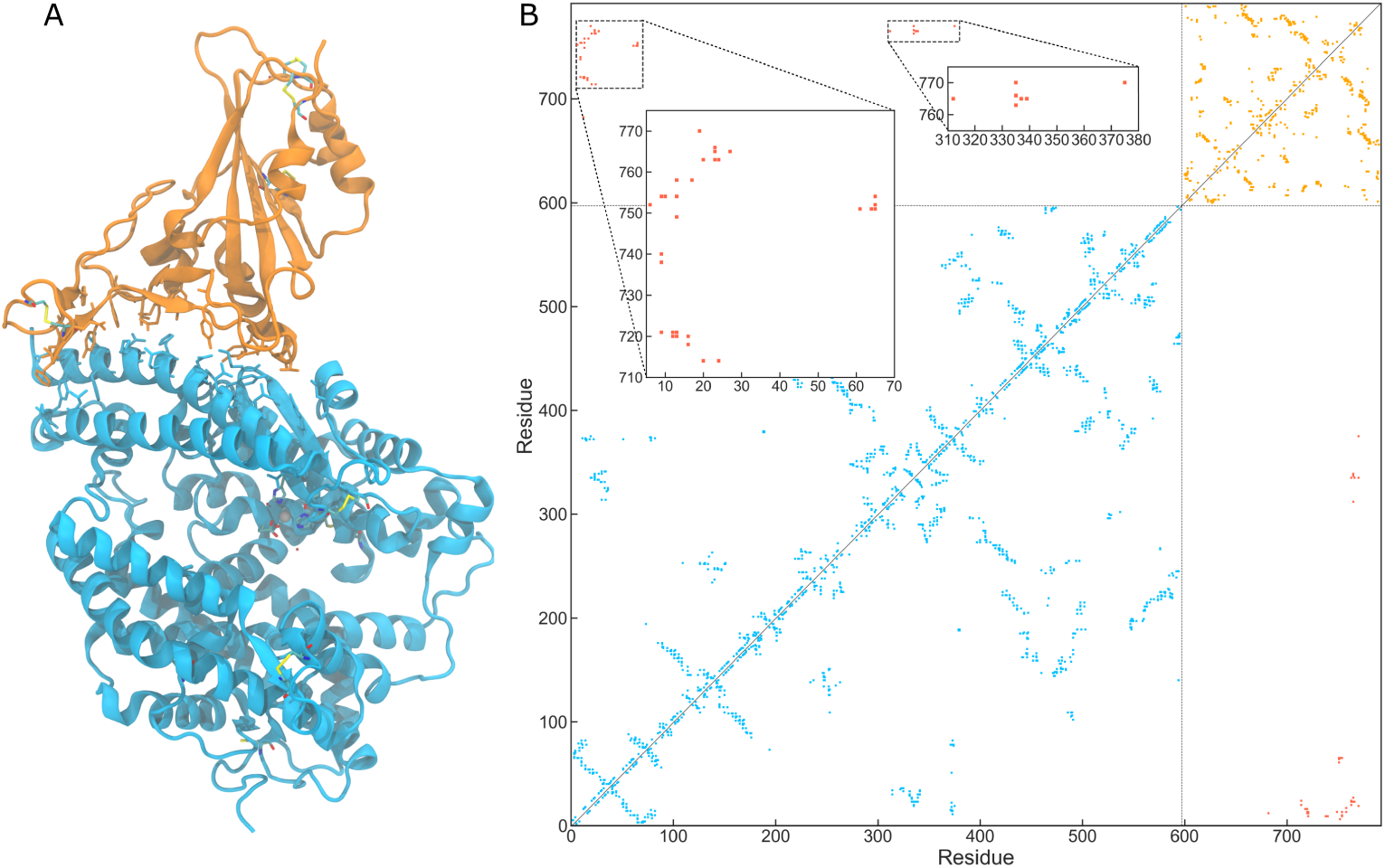
Three dimensional structure (A) and contact map (B) of the ACE2:RBD complex. ACE2 is shown in cyan and the RBD in orange. We show atomic detail for residues forming intermolecular contacts, cysteines and Zn coordinating residues in ACE2. Intra-molecular contacts for each protein in (B) are shown in the same colours as in (A), while intermolecular contacts are shown in red. The insets highlight the regions with most intermolecular contacts.

In order to better understand viral infection and design ways to hijack it, it is important to understand the mechanism of binding between the spike protein and ACE2. This information is not readily available from the static X-ray and cryo-EM experimental structures. Molecular dynamics (MD) simulations are ideally posed to provide this picture. However, simulating protein-protein binding using atomistic MD even for small proteins is exceedingly time-consuming even for much smaller systems than the RBD:ACE2 complex [12, 13]. Alternatively, one may resort to coarsegrained structure-based (Gō) models, whose potential energy minimum is located by construction in the reference experimental structure [14]. In this way, these models are consistent with the central prescription of a minimally frustrated, funnelled energy landscape of protein folding theory [15]. Despite their simplicity, structure based models provide a description of folding mechanisms that is consistent with that of atomistic MD in explicit solvent [16], and have resulted in quantitative agreement with experiment [17, 18]. In the past, Gō models have been applied successfully to the study of protein-protein binding for complexes involving both folded and disordered proteins [19, 20, 21].

In this work, we use the structure-based simulation model by Karanicolas and Brooks [22], to characterize the free energy landscape for RBD binding to the ACE2 receptor. We calibrate the model in order to recover the experimental binding affinity and subsequently explore the emerging features. Additionally, from equilibrium runs we describe the binding reaction paths. We find considerable disorder in the binding of RBD to ACE2 and that the binding mechanism is dominated by the interactions involving the receptor binding ridge.

## 2. Methods

### 2.1. Coarse-grained molecular simulations

We use the structure-based simulation model by Karanicolas and Brooks, which is coarse-grained to the C^*α*^ level [22]. The potential energy function in the model is defined as a sum of terms for bonds, angles, dihedrals and non-bonded interactions

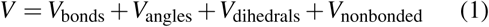

The first two terms in this expression are harmonic potentials centered in the value of the bond or angle in the reference (experimental) conformation. The dihedral term is based on statistical propensities for torsion angles in the PDB. The non-bonded term is attractive, with a modified Lennard-Jones potential,

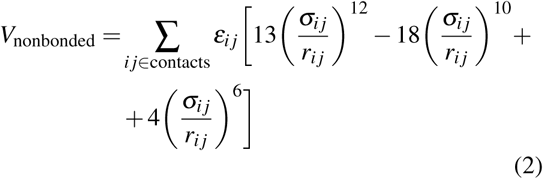

for pairs of beads corresponding to amino acids that are in contact in the native structure (have atoms within 4.5 Å). For every other pair of beads, a repulsive core is defined [22].

We built models using a recently determined 2.45 Å resolution X-ray structure of the ACE2:RBD complex (with PDB ID: 6m0j [5], see Figure 1A). Although we take into consideration both the disulfide bonds and Zn(II) interactions in our simulations we do not consider glycosilation effects. Among the 22 predicted N-glycosylation sites in the spike protein of SARS-CoV-2 [23], only the highly conserved glycosylation site N343 is in the RBD, and this is located far from the interaction region [11]. It is also relevant to note that a second glycosylation site in the RBD of the SARS-CoV spike (N357) is lost in SARS-CoV-2 [23]. The energy terms in the model were calibrated carefully in order to recover a value of *K*_D_ at 300 K in the experimental range (i.e. tens of nM, see Supplementary Information Fig. S1).

We run replica exchange simulations (REMD) using 32 replicas at temperatures ranging between 285 and 378 K, evenly spaced every 3 K. We run a total of 2 *µ*s for each of the replicas, using a Langevin dynamics integrator with a 10 fs time-step and a friction constant of 0.2 ps^*-*1^. Additionally, we run short equilibrium trajectories starting at randomly displaced and rotated RBD conformations within the unbound state. All the simulations were run using the Gromacs software package [24].

### 2.2. Analysis methods

We project the simulation data on the intramolecular fraction of native contacts, *Q*, defined as the average degree of contact formation

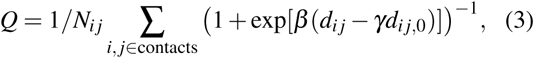

where *d*_*i j*_ and *d*_*i j*,0_ are the distances between beads *i* and *j* in an instantaneous and the reference conformations, and the sum runs over pairs of residues forming native contacts [25]. As a binding coordinate we use the intermolecular *d*_RMS_ defined as [26],

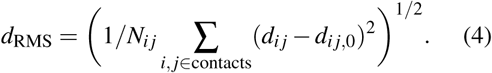

We combine the data from different runs to derive free energy surface using WHAM [27]. The dissociation constant *K*_D_ was calculated as

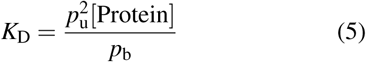

where *p*_b_ and *p*_u_ are the populations of the bound and unbound states, and the protein concentration is straightforwardly derived from the simulation box volume as before [28].

## 3. Results and Discussion

In order to obtain the free energy landscape of RBD binding to ACE2 we first run REMD simulations that enable exhaustive exploration of conformational space. The resulting 2 *µ*s of simulation data at each temperature were projected on an order parameter for binding, the logarithm of the *d*_RMS_, that collapses the unbound state into a narrow well and expands the bound state (see Methods). The simulations are well converged, as shown by cumulative averages that do not change after ∼1 *µ*s at each temperature (see Supplementary Information Fig. S2).

In Figure 2A we show potentials of mean force derived using WHAM [27]. The free energy surface suggests that the RBD:ACE2 complex formation occurs in an all-or-none way, with a dominant free energy barrier between the bound and unbound states. Despite the absence of stable intermediates, the bound state has a small intra-state barrier. The presence of two sub-basins in the bound state with average values of log(*d*_RMS_) of -0.45 and -0.75 is also apparent in the two dimensional free energy landscape we show in Figure 2B, including also a folding coordinate for the RBD, the fraction of native contacts, *Q*_RBD_.

**Figure 2:**
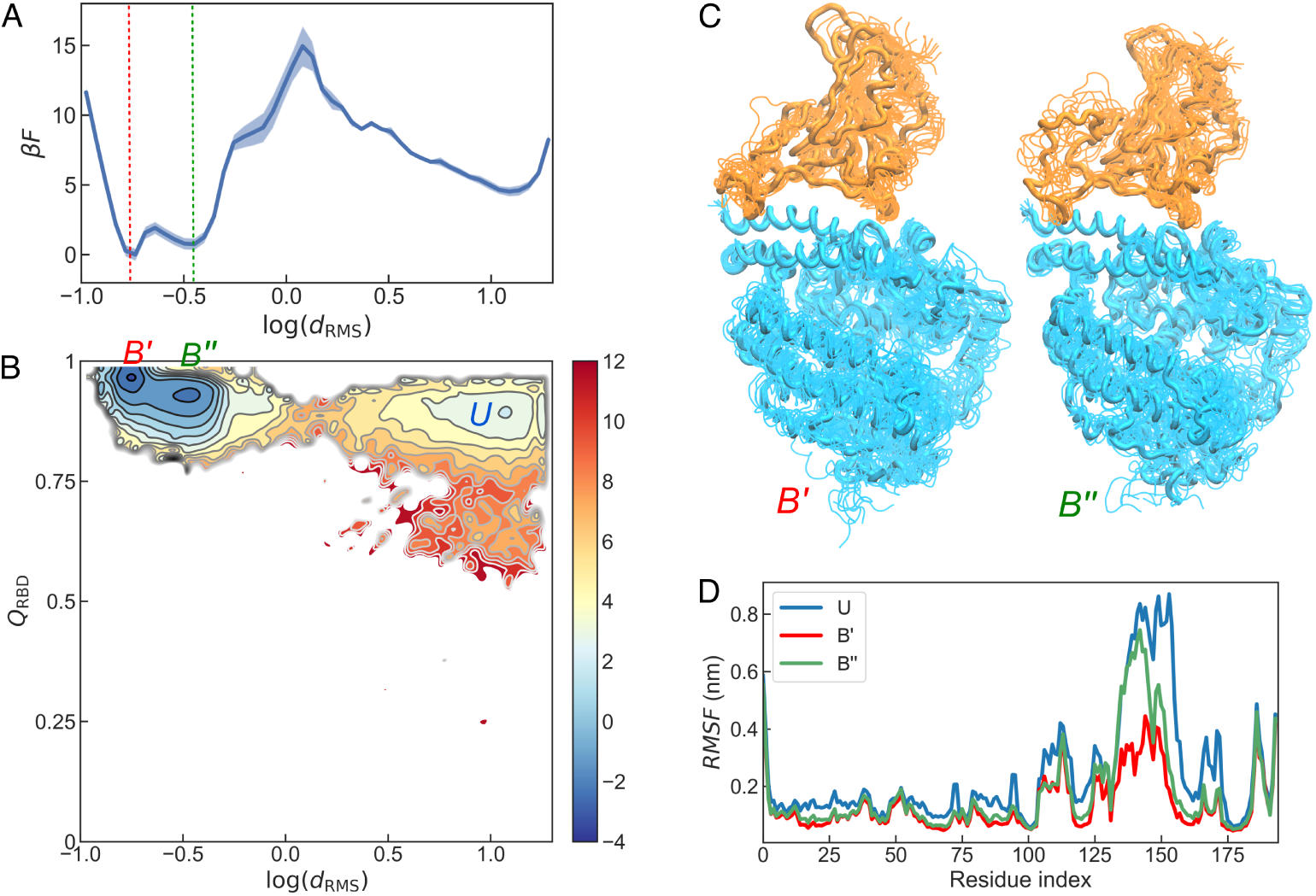
(A, B) Potentials of mean force for the binding of SBD to ACE2 for the projection on log(*d*_RMS_) and both the fraction of intra-RBD contacts *Q*_RBD_ and log(*d*_RMS_). B’ and B” are the fully bound and fluctuating sub-basins within the bound state and U marks the unbound state. Free energies are in units of *kT*. (C) Representative snapshots for the B’ and B” states. (D) *RMSF* for RBD in the different states as a function of the residue index.

Hence, the free energy landscapes suggest some degree of heterogeneity in RBD even in the bound state. In Figure 2C we show snapshots of conformations within each of the bound state sub-basins, derived from the REMD simulation data at 300 K. The lowest log(*d*_RMS_) form (which we term B’) very much matches the experimental structure, while in the higher log(*d*_RMS_) form (B”) there are considerable conformational fluctuations that seem mainly localized in the region termed receptor binding ridge [7, 11], despite the disulfide bond that barely holds this region folded.

To characterize the residual disorder in the bound state we calculate the root mean square fluctuations (*RMSF*) for the RBD estimated from snapshots of both substates and the unbound state (see Figure 2D). As anticipated, fluctuations are primarily localized in the ridge region, both in the unbound and in the bound form of the protein, although in the former they extend a bit longer to the receptor binding region. We see that upon binding, fluctuations decrease considerably in the RBD but they remain larger than average in the receptor binding ridge. Although Gō models underestimate the cooperativity of conformational transitions, and may hence overemphasize the fluctuations, there is external support for this prediction. On one hand,the ridge region could not be properly resolved in recent cryo-EM structures of the spike protein [4]. Also, structural disorder is predicted to appear around residue 150 of RBD from multiple servers [29]. Additionally, the location of the fluctuations of the coarse-grained simulations is qualitatively consistent with atomistic MD [30]. Finally, we have built an elastic network model from the experimental structure of the RBD using the ProDy software package [31], which also gives consistent results (see Supplementary Figure S4). All of this evidence gives further confidence on the outcome of our simulations.

In order to recover a dynamic picture of the binding process we have run a hundred independent simulation runs starting from randomly oriented conformations in the unbound state, resulting in 22 binding events. This allows to understand the importance of the properties of the underlying energy landscape in the binding process, in particular the relevance of disorder in RBD, and propose a binding mechanism. In Figure 3 we show the projection on relevant progress coordinates of the simulation data for one of the binding events (two more are shown in the Supplementary Information Fig. S3). Again, we monitor the log(*d*_RMS_) as a progress coordinate for binding, but now considering either all contacts or different subsets of contacts separately. In particular, we discern between contacts formed by the ridge residues and those in the remaining part of the extensive RBM. In the overwhelming majority of the events we have observed, binding occurs first via the disordered region, which establishes the first few contacts, resulting in a decrease of log(*d*_RMS_). Then, the remaining part of the RBD rests on the ACE2 binding site, until the protein fully commits to the bound form. Representative snapshots help visualize how the extended, disordered part of the protein more efficiently searches for its binding partner, possibly due to a slightly increased capture radius (Figure 3).

**Figure 3:**
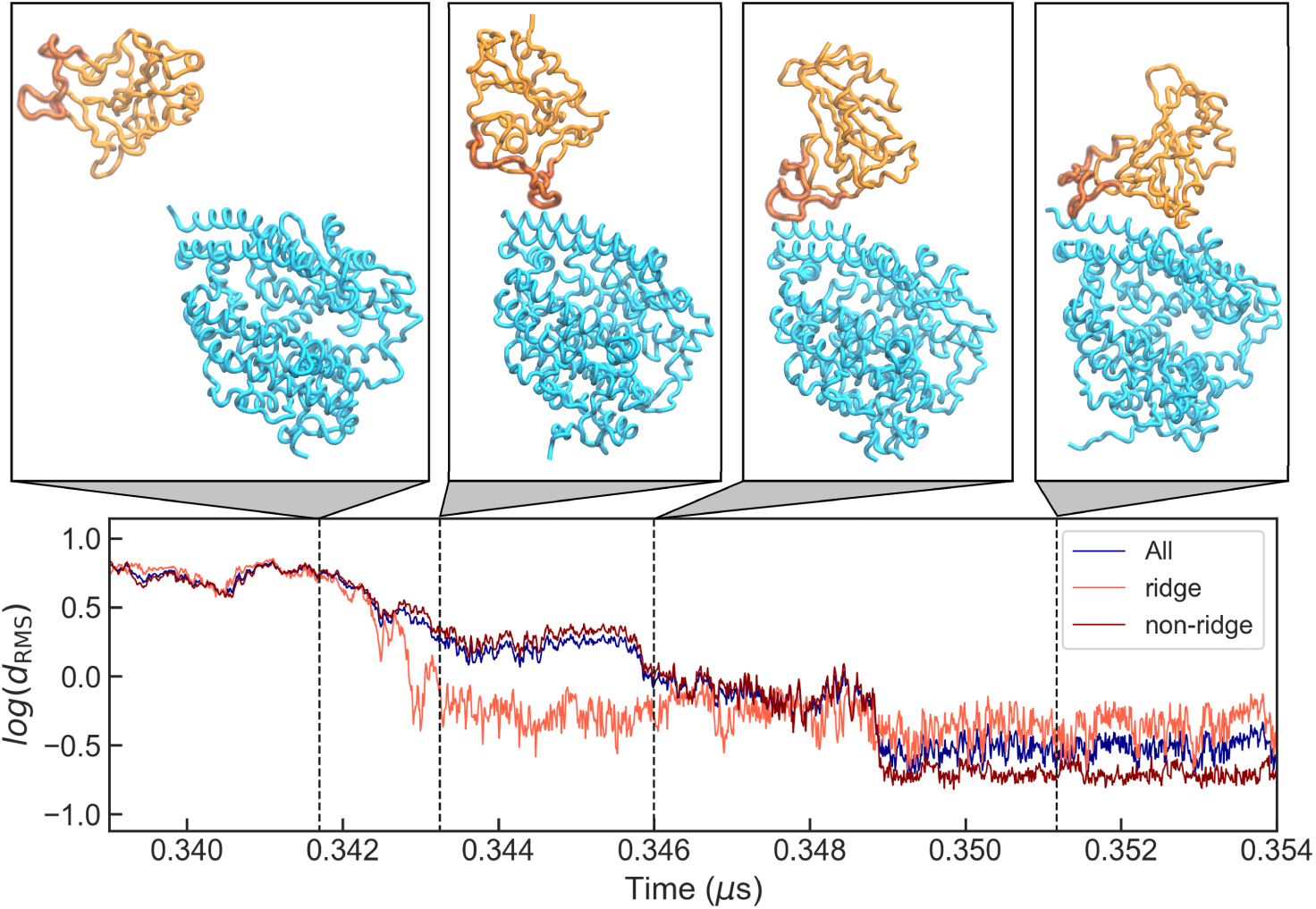
Representative transition path for a binding transition between RBD and ACE2. We show the projection of the simulation data on the log(*d*_RMS_) for all contacts (dark blue) or different subsets of interprotein contacts, either involving the ridge or not (light and dark red, respectively). Snapshots at different time-points are shown at the top. Colours as in Fig. 1. The thicker region in the RBD corresponds to the ridge.

Our results provide, to our knowledge, the first molecular description of the binding process of the RBD to the ACE2 receptor which is an essential first step in the infection by COVID-19. One of the key insights is the role of structural disorder in the binding ridge of the RBD. In the last years, considerable attention has been dedicated to the role that disordered loops may have in conformational dynamics and allostery [32]. The preferential binding through the disordered region is also reminiscent of the so-called “fly-casting” mechanism that has been invoked to explain the kinetic advantage of intrinsically disordered proteins for binding [33]. In the particular case of viral proteins, structural disorder has recently been highlighted as a prevalent feature [34], suggesting that small disordered regions may provide an evolutionary advantage.

There are several potential limitations in our work due to the coarse-grained nature of the model and the simplicity in the description of the interactions. First, the simulation model does not consider explicitly longrange charged interactions, although pairwise energy terms are residue-specific [22]. This prevents accounting for electrostatic steering effects that may speed up the binding process [35]. Additionally, we do not incorporate the glycosilation effects, although experimental structures show that glycosilated sites in ACE2 and RBD are far from the binding interface and hence likely to be small.

We hope the insight from our coarse grained simulations will be helpful in future drug discovery work either for antivirals that exploit the binding properties of RBD:ACE2 or for vaccine development. One possibility would be to try to tune the disorder in the RBD, distorting the balance between conformational entropy loss and enthalpic gain. Alternatively, fragments of ACE2 involving the regions of the protein forming most of the native contacts may be tested as potential inhibitors of the virus.

## Acknowledgments

DDS receives financial support from the grants PGC2018-099321-B-I0 and RYC-2016-19590 from the Spanish Ministry of Science, Research and Universities (MINECO/FEDER) and the Basque Government through grant IT588-13. The Spanish Ministry of Science, Research and Universities also supports JAG and RPJ through grants BIO2016-74875-P and BIO2016-77390-R, respectively. DDS wishes to thank Robert B. Best for generously sharing computational tools and the staff at the DIPC Supercomputing Center for technical support. DDS also acknowledges María M. Caffarel, for helpful discussions and Carmen and Lucía de Sancho for key insights in the structure of the complex.

