## Supplementary Information for "Coarse-grained molecular simulations of the binding of the SARS-CoV-2 spike protein RBD to the ACE2 receptor"

### Supplementary Methods

#### Gaussian Network Model of RBD

In order to explore the global biomolecular dynamics of the RBD, we have produced an elastic network model of RBD using the software package ProDy [1]. As the only input we use the molecular coordinates of the same structure of the RBD of the spike protein used to generate the simulation models [2]. In Figure S4 we report the fluctuations as a function of the residue number, which give an indication of the most dynamic regions of the protein.

### Supplementary Figures

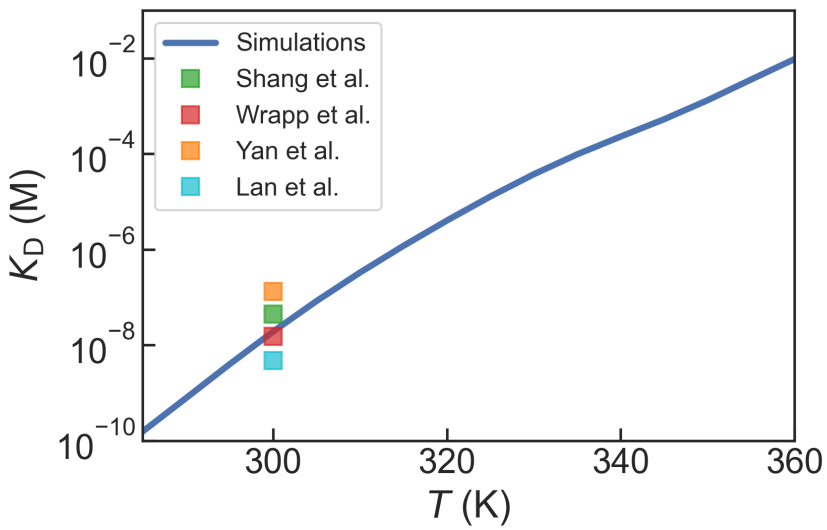


Figure S1: Temperature dependence of the dissociation constant in the calibrated molecular simulation model. We show the values from different experimental groups as squares[2–5].

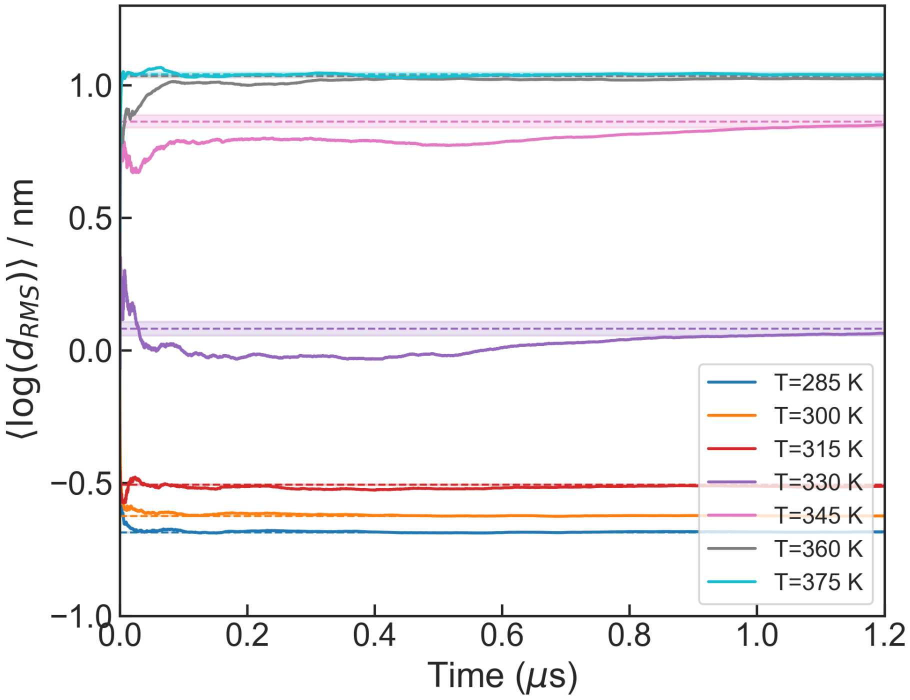


Figure S2: Running averages of the $\log(d_{\mathrm{RMS}})$ as a function of time, at multiple temperatures. Dashed lines and bands mark averages and shaded bands.

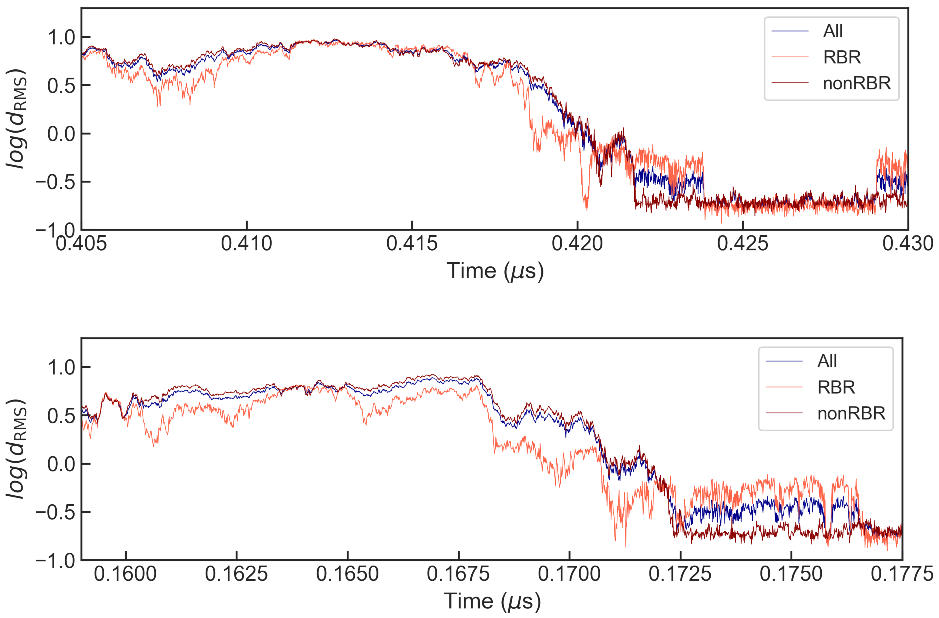


Figure S3: Transition paths for two additional binding events.

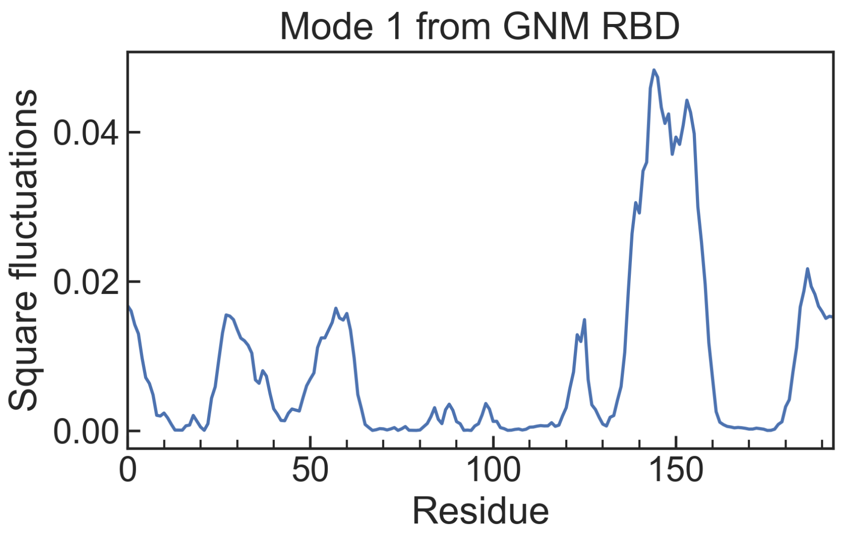


Figure S4: Squared fluctuations for the first mode from the gaussian network model
